# Increased alcohol self-administration following exposure to the predator odor TMT in high stress-reactive female rats

**DOI:** 10.1101/2020.07.17.208561

**Authors:** Laura C. Ornelas, Ryan E. Tyler, Preethi Irukulapati, Sudheesha Paladugu, Joyce Besheer

**Affiliations:** Bowles Center for Alcohol Studies, University of North Carolina at Chapel Hill, Chapel Hill, North Carolina 27599; Neuroscience Curriculum, University of North Carolina at Chapel Hill, Chapel Hill, North Carolina 27599; Department of Psychiatry, University of North Carolina at Chapel Hill, Chapel Hill, North Carolina 27599

**Keywords:** Predator odor stress, TMT, individual differences, alcohol, sex differences, post-traumatic stress disorder, alcohol use disorder

## Abstract

Post-traumatic stress disorder (PTSD) and alcohol use disorder (AUD) are highly comorbid. Additionally, individual differences in response to stress suggest resilient and susceptible populations. The current study exposed male and female Long Evans rats to the synthetically produced predator odor 2,5-dihydro-2,4,5-trimethylthiazoline (TMT) to examine individual differences in stress-reactive behaviors (digging and immobility) and whether these differences could predict lasting consequences of TMT and increases in alcohol drinking. Male and female Long Evans rats were trained on operant alcohol self-administration. After 9 sessions, rats underwent exposure to TMT or water (Control) in a distinct context. 6 days after TMT exposure, rats underwent re-exposure to the TMT-paired context (without TMT), and a series of behavioral assessments (acoustic startle, zero maze, light/dark box), after which rats resumed alcohol self-administration. Rats were divided into two TMT-subgroups using a ratio of digging and immobility behavior during TMT exposure: TMT-subgroup 1 (low digging/immobility ratio) and TMT-subgroup 2 (high digging/immobility ratio). Digging/immobility ratio scores predicted elevated corticosterone levels during TMT exposure and reactivity during context re-exposure in males and females (TMT-subgroup 2), as well as elevated corticosterone levels after context re-exposure and hyperarousal behavior in females (TMT-subgroup 1). Furthermore, TMT stress reactivity predicted increases in alcohol self-administration, specifically in females. These data show that stress-reactivity can predict lasting behavioral changes which may lead to a better understanding of increases in alcohol drinking following stress in females and that individual differences in stress-reactive behaviors using TMT may be helpful to understand resilience/susceptibility to the lasting consequences of stress.

**Highlights:**

- Exposure to the predator odor TMT produces distinct behavioral phenotypes in male and female rats
- Male and female high stress reactive rats show enhanced reactivity to the TMT-paired context
- Stress-reactivity during TMT predicts increases in alcohol self-administration, in females
- Stress-reactivity may help to understand resilience/susceptibility and impact on alcohol drinking

## 1. Introduction

Post-traumatic stress disorder (PTSD) is an anxiety, trauma-and stressor-related disorder that manifests after exposure to a traumatic event. Diagnostic criteria include exposure to a traumatic stressor, intrusive symptoms, avoidance or negative alterations in cognition and mood, and alterations in arousal and reactivity [1]. Lifetime prevalence of PTSD in the United States is approximately 8.3% for adults, with females (12.8%) being more likely to have a lifetime prevalence of PTSD compared to males (5.7%) [2, 3]. Furthermore, it is well known that PTSD is highly comorbid with alcohol use disorder (AUD) [4, 5]. According to the National Comorbidity Study [4], 26.2% of women and 10.3% of men in a general population with alcohol dependence have met the criteria for PTSD. Some individuals with PTSD consume alcohol as an attempt to alleviate symptoms, which can increase the risk of developing a drinking problem [6]. In addition, experiences of trauma despite the diagnosis of PTSD, can induce high levels of alcohol craving and lead to an increase in consumption [7].

Not all individuals exposed to trauma develop PTSD. Approximately 5-10% of individuals develop PTSD after experiencing a traumatic stressor [2]. Studies examining stress resilience and susceptibility in humans show one’s ability to adapt to stressful encounters are key factors that can predict resilient outcomes to stress [8]. Well-adapted behavioral responses such as active coping mechanisms [9] and cognitive reappraisal strategies [10] prompt a level of resiliency that can protect against developing PTSD [11, 12]. Therefore, it is necessary to identify novel animal models to target behavioral and neurobiological mechanisms associated with individual variability in response to stress and trauma to better understand differences in vulnerability to developing PTSD.

Animal models have become increasingly important in stress research to examine behaviors that can inform our understanding of clinical PTSD symptoms. These models can be used to investigate the relationship between increased alcohol consumption and stress including examining the relationship between individual differences in symptom profiles of traumatic stress and excessive alcohol drinking [13, 14]. Specifically, predator odor exposure including soiled cat litter [15] and bobcat urine [16] have been shown to produce escalations in alcohol drinking in rats, while dirty rat bedding has been shown to increase alcohol consumption in mice [17]. Additionally, exposure to 2,3,5-trimethyl-3-thiazoline (TMT; an extract of fox feces) has been shown to produce alcohol reinstatement (e.g., relapse-like behavior) in mice [18]. Animal models can also be used to examine relevant individual variability in responses to stress [19-24], including stress resilience. Previous studies have utilized a variety of methods to classify animals into specific phenotype groups to examine individual variability following the stress exposure. For example, classification can be based on avoidance behavior during re-exposure to a bobcat urine paired context [16, 25], behavior during an elevated-plus maze and acoustic startle response following TMT exposure [16, 25], and anxiety-like behavior in an elevated plus maze and context avoidance behavior [23, 24]. These characterization methods focus on grouping rats based on behavioral changes that occur after exposure to stress. The focus of the current study was to determine whether quantifying behavior during the stressor exposure could provide an index of stress-reactivity that could be used to predict long-term consequences of stress including context reactivity, anxiety-like behavior, hyperarousal behavior, and increases in alcohol self-administration in male and female rats.

The current study uses TMT exposure as the stressor because it activates a hardwired “learned-independent system” shown to induce innate fear and defensive behaviors [26]. An important advantage of using predator odor exposure, including TMT, as a stressor, is the ability to measure stress-reactive behaviors during stressor exposure, including defensive digging and immobility [27], which can be used as an index of stress-reactivity. Digging is a species-and strain-dependent behavior that has been implemented in rodent models of PTSD [27, 28]. It is interpreted as a proactive response to stress [29-31], such that it reflects an active coping response or fear-related behavior [32] and predator-stress responsiveness [33]. Freezing behavior is a well-characterized fear-like behavior that animals have been shown to engage in during TMT exposure [27, 34-36]. The present study sought to examine individual differences in stress-reactive behaviors (digging and immobility) during TMT exposure in male and female Long Evans rats to determine whether these individual differences could predict subsequent increases in context reactivity, hyperarousal and anxiety-like behavior and alcohol self-administration. Rats were divided into two TMT-subgroups using a ratio of defensive digging and immobility behavior during TMT exposure (digging/immobility ratio score) to examine the relative proportion of digging behavior to immobility behavior. Digging and immobility during TMT were specifically chosen as the measures to calculate ratio scores because they represent two distinctly different types of stress-induced behavioral coping strategies [29, 31, 37, 38] (digging = active, immobility = passive) that can be used to reflect individual differences in stress responsivity. We hypothesized that higher digging/immobility ratio scores, indicative of high levels of digging and low levels of immobility, would predict increases in reactivity to the TMT-paired context upon re-exposure, anxiety-like and hyperarousal behaviors, and subsequent increases in alcohol self-administration.

The current study supports several important findings including 1) male and female rats show individual variability in the engagement of stress-reactive behaviors during TMT, specifically immobility behavior and 2) stress reactivity during TMT exposure can be an important predictor of subsequent alcohol self-administration, specifically in female rats. These lasting consequences of exposure to the synthetically produced predator odor TMT can provide a better understanding of stress-induced increases in alcohol drinking, as well as how individual differences and heterogeneity in stress-reactive behaviors using TMT may be helpful in understanding individual resilience/susceptibility to stress and its lasting consequences.

## 2. Materials and Methods

### 2.1. Subjects

Male and female young adult (arrived at 7 weeks old) Long Evans rats (n=64) were used for these experiments. Animals were double housed by sex in ventilated cages (Tecniplast, West Chester, PA) upon arrival to the vivarium with ad libitum food and water. For at least 1 week prior to the start of alcohol self-administration training, all rats were single housed and remained single-housed for the duration of the experiment. Rats were maintained in a temperature and humidity-controlled colony with a 12-hour light/dark cycle (lights on at 07:00). All experiments were conducted during the light cycle. Animals were handled for five days prior to the start of the experiment. Animals were under continuous care and monitoring by veterinary staff from the Division of Comparative Medicine at UNC-Chapel Hill. All procedures were conducted in accordance with the NIH Guide to Care and Use of Laboratory Animals and institutional guidelines.

### 2.2. Self-Administration Apparatus

Self-administration chambers (31 × 32 × 24 cm; Med Associates Inc.; St. Albans, VT) were individually located within standard-attenuating cubicles equipped with an exhaust fan that provided both ventilation and masking of external sounds. Chambers were fitted with a retractable lever on the left and right walls and a white cue light was centered 7-cm above each lever with a liquid receptacle centered on each wall. Lever responses on the left lever (i.e. active lever) activated a syringe pump (Med Associates) that delivered 0.1 ml of solution into the receptacle during a 1.66-s period. The white cue light and tone located above the active lever were activated during pump activation. Responses on the right lever (i.e. inactive lever) had no programmed consequence. The chambers also had infrared photobeams which divided the floor of the chamber into 4 zones to record general locomotor activity throughout each session.

### 2.3. Alcohol Self-Administration Training

The experimental timeline is illustrated in Figure 1. Rats were trained on operant alcohol self-administration training using a sucrose-fading procedure in which alcohol was gradually added to the 10% (w/v) sucrose solution, similar to [39]. The exact order of daily exposures was as follows: 2% (v/v) alcohol/10% (w/v) sucrose (2A/10S), 5A/10S, 10A/10S, 10A/5S, 15A/5S, 15A/2S, 20A/2S, 20A with one day at each concentration except for 20A which occurred for two days after which the rats remained on 15A as the reinforcer for the remainder of the study. Self-administration sessions (30 minutes) took place 5 days per week (M-F) with the active lever on a fixed ratio 2 schedule (FR2) of reinforcement such that every second response resulted in delivery of alcohol.

**Figure 1.**
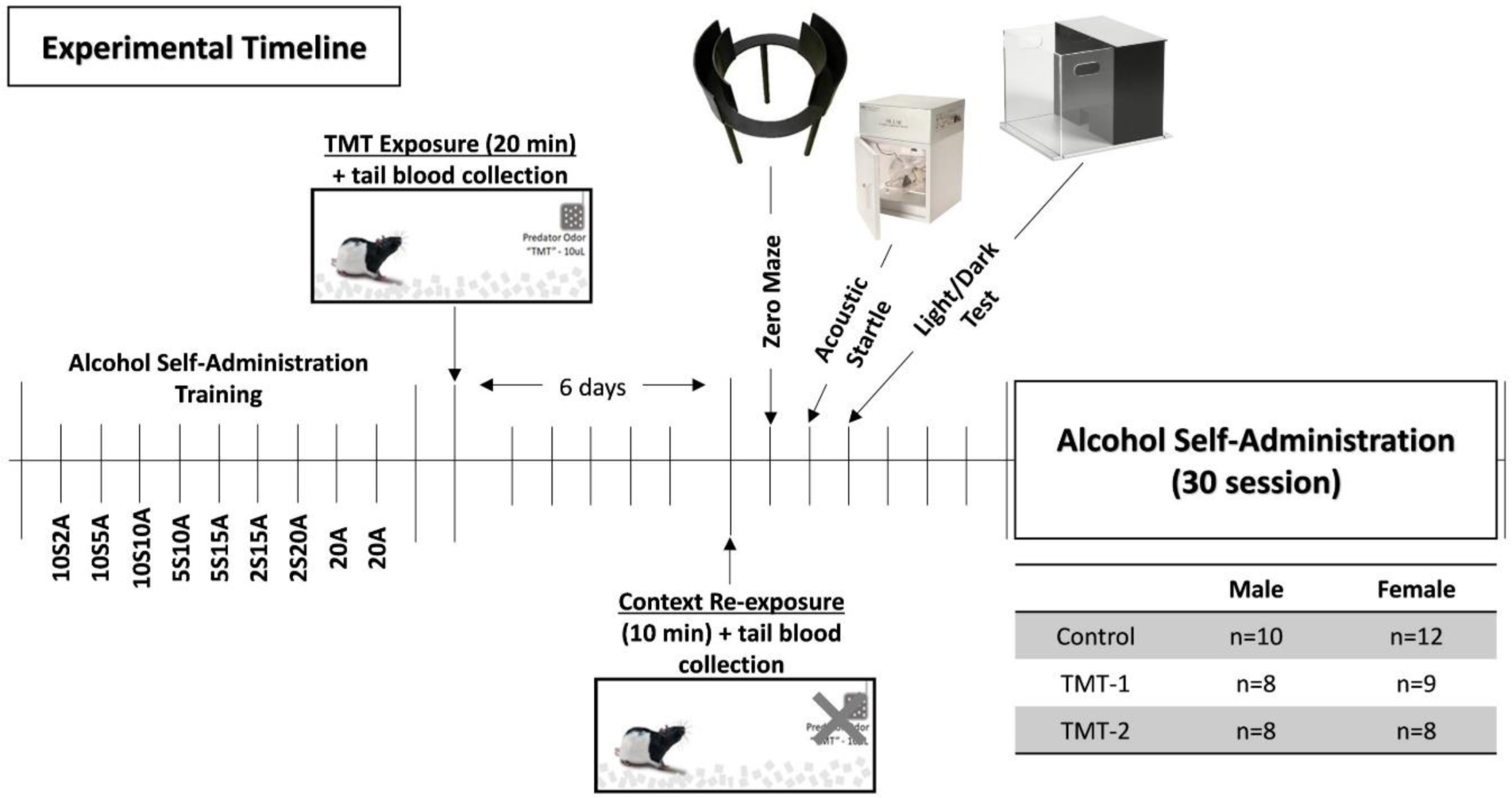
Experimental Timeline. Male and female Long-Evans rats were trained on alcohol self-administration for 9 days. 24 hr post last self-administration training day, male and female rats were exposed to water or TMT. 6 days later, rats were re-exposed to the initial TMT-paired contextual environment in the absence of TMT. 24 hr later rats underwent testing for anxiety-like behavior (zero maze, hyperarousal (acoustic startle response, ASR) and anxiety-like behavior as measured by approach/avoidance behavior (light/dark test and zero maze). Three days after behavioral tests, rats returned to alcohol self-administration for 30 sessions (four weeks). The sample size of each treatment group for male and female rats is also included.

### 2.4. Predator Odor Exposure and Context Re-exposure

Approximately 24 hr after nine days of self-administration training, rats underwent predator odor exposure. Rats were transferred from the home cage to Plexiglas exposure chambers (45.72 × 17.78 × 21.59 cm; UNC Instrument Shop, Chapel Hill, NC, USA). The length of the back wall of the chambers was opaque white with two opaque black side walls and a clear front wall to allow for video recordings. A metal basket (17.8 cm above the floor) was hung on the right-side wall. This basket held a piece of filter paper on which was placed 10 μl of TMT or water (for controls) so that the filter paper was inaccessible to the rat. Approximately 600 mL of white bedding (Shepherds ALPHA-dri) was added to the bottom of the exposure chamber prior to the animal being placed in the chamber. After the rat was placed into the chamber, a clear Plexiglas top was slid and secured into place. The exposure session was 15 min in duration and recorded by a video camera for later analysis using ANY-maze ™ Video Tracking System (Stoelting Co. Wood Dale, IL, USA). After completion of the exposure, fecal boli was counted for each rat as a measure of physiological responses to TMT. Following the exposure, rats were immediately transferred to a separate procedure room where tail blood was collected for later analysis of plasma corticosterone (CORT levels. Following the completion of blood collection, each rat was returned to the homecage. Blood was collected into heparinized tubes and centrifuged at 4°C for 5 minutes at 2000 rcf. Approximately 20-30 µL of plasma supernatant was collected and stored at −80°C until analysis. Plasma samples (5 µL) were analyzed for corticosterone levels in duplicate using colorimetric EIA kit (ArborAssays, Ann Arbor, MI) per the manufacture instructions.

One week following predator odor exposure, reactivity to the TMT-paired context was examined. Animals were transferred from their home cage and placed in the context in which they had been previously exposed to water or TMT for 10 min (no TMT present). Sessions were recorded by a video camera and analyzed using ANY-maze ™ Video Tracking System (Stoelting Co. Wood Dale, IL, USA). Following the 10 min context re-exposure, rats were immediately transferred to a separate procedure room where tail blood was collected for later analysis of plasma corticosterone levels. Following the completion of blood collection, each rat was returned to the homecage.

### 2.5. Elevated Zero Maze

24 hr after the context re-exposure, rats underwent testing in an elevated zero maze to assess anxiety-like behavior. The zero maze was composed of a circular platform with a diameter of 99 cm raised above the floor to a height of approximately 70 cm and divided equally into four quadrants. Two enclosed quadrants contain two walls with one back wall [33.02 cm (H)] and front wall [25.4 in (H)]. The exposed quadrants are located on opposite sides of the circular platform and are bordered with approximately 5 mm rim in order to prevent the rat from falling off the circular platform. Rats were placed in one open quadrant at the start of the test for 5 min in the zero maze before being placed back into their home cage.

### 2.6. Acoustic Startle Response

On the following day, rats underwent acoustic startle response testing to assess changes in arousal response using an acoustic startle response system (S-R Lab; San Diego Instruments, San Diego, CA). Rats were placed in a cylinder Plexiglas animal enclosure located within a sound-attenuating test chamber that included an exhaust fan, a sound source, and an internal light that was turned off during the test. At the start of each test, rats underwent a 5-min habituation period during which 60 dB of background white noise was present. The background noise was present during the entire test session. The test session consisted of 30 trials of a 100 ms burst of a 110 dB startle. Each trial was separated by a 30 to 45-s randomized intertrial interval. Startle amplitude was measured with a high-accuracy accelerometer mounted under the animal enclosure and analyzed with SR-Lab software.

### 2.7. Light/Dark Test

The day after the acoustic startle test (72 hr after the context re-exposure test), animals underwent testing for anxiety-like behavior as measured by approach/avoidance behavior in the light/dark chamber. A dark box insert (44.4 × 22.9 × 30.5 cm) was placed in the left side of an open field chamber (150 lux) (23.31 × 27.31 × 20.32 cm; Med Associates Inc.; St. Albans, VT) to divide the chamber into a dark and light side. The chamber was located within standard-attenuating cubicles equipped with an exhaust fan that provided both ventilation and masking of external sounds. Time and distance spent on each side of the chamber was measured with 4 parallel beams across the chamber floor. Animals were transported into the testing room in the home cage at least 20 min prior to the start of the 5 min test and each rat was placed in the light side facing the posterior wall.

### 2.8. Alcohol Self-Administration

Rats resumed alcohol self-administration 4 days after completion of the light/dark test. 30-min self-administration sessions took place 5 days per week (M-F) for 20 sessions (15% v/v, alcohol). After these 20 sessions, self-administration sessions occurred 3 days per week (MWF) for 4 weeks for the remainder of the study.

### 2.9. Data Analyses

#### 2.9.1. TMT and Context Re-exposure

TMT Subgroup Classification. Male and female rats exposed to TMT were grouped into two TMT-subgroups by a digging/immobility score based on their behavior during the TMT exposure. This score was calculated by dividing the total time spent digging by the total time spent immobile. A median split of the scores was performed to divide the range of ratio scores into two TMT-subgroups for each sex; lower ratio scores were classified as TMT-subgroup 1 and higher ratio scores were classified as TMT-subgroup 2 in both male and female rats. Using ANY-maze, the length of the rectangular TMT exposure chamber was divided into two compartments for analysis (TMT side and non-TMT side). Immobility was operationally defined as lack of movement for more than 2 seconds as assessed using ANY-maze software. Therefore, immobility likely captures both inactivity and freezing, which is characteristic of a fear response in rodents and observed during TMT exposure [34]. One-way RM analysis of variance (ANOVA) tests were used to compare differences in control and TMT-subgroups in male and female rats. A video recording error during the TMT exposure resulted in two female rats in the TMT group having to be excluded from the entire study. Cumulative time spent digging, immobility and time spent on TMT side was calculated across 10 min of exposure rather than the total 15 min exposure. One male TMT rat determined to be a statistical outlier based on digging behavior during TMT (greater than 2 standard deviations from the mean) was excluded from the study. Analysis of stress-reactive behaviors across time was examined by two-way RM ANOVA with TMT exposure as a between-subjects factor and time as a within-subjects factor. Tukey multiple comparisons tests were used to follow up significant main effects of groups and interactions. All data are reported as mean ± SEM. Significance was set at p≤0.05.

#### 2.9.2. Light/Dark Test, Acoustic Startle Response and Zero Maze Test

The % time in light compartment for the light/dark test, average startle amplitude for acoustic startle response and % open arm time for the zero maze were analyzed by a one-way RM ANOVA. Tukey multiple comparisons tests was used to analyze significant main effects of groups. All data are reported as mean ± SEM. Significance was set at p≤0.05.

#### 2.9.3. Self-Administration

For alcohol self-administration, alcohol lever responses and alcohol intake (g/kg) are presented as 3-session averages. Alcohol intake was estimated from reinforcers received relative to body weight (kg). These data were analyzed by a two-way RM-ANOVA. Tukey multiple comparisons tests was used to analyze significant main effects of groups and interactions. General locomotor activity during the session was measured and total beam breaks across the session was divided by the session length (30 min) to determine locomotor rate (beam breaks/min). A baseline criterion was set such that rats that had below an average of at least 0.30 g/kg between the two 20A baseline training days prior to TMT exposure were excluded from the study. Two control-male rats, three TMT-male rats and one TMT-female did not meet this criterion. All data are reported as mean ± SEM. Significance was set at p≤0.05.

## 3. Results

Prior to TMT exposure, there were no differences in alcohol self-administration as measured by lever responses across training days, total alcohol drinking history (Table 1, *p* > 0.05), locomotor activity or inactive lever responses (*p* > 0.05) between TMT-subgroups and controls in male or female rats.

**Table 1.** Alcohol lever responses and alcohol intake (g/kg) from sucrose fading phase during self-administration training

|  |  | Alcohol Lever Responses |  |  |  |  |  |  |  |  |
| --- | --- | --- | --- | --- | --- | --- | --- | --- | --- | --- |
| Group |  | 10S2E | 10S5E | 10S10E | 5S10E | 5S15E | 2S15E | 2S20E | 20E | 20E |
| Male | Control | 81.5 ± 22.1 | 85.6 ± 18.3 | 106.7 ± 19.8 | 96.5 ± 15.8 | 69.6 ± 14.0 | 33.4 ± 6.0 | 50.2 ± 8.1 | 25.9 ± 4.7 | 30.0 ± 5.2 |
|  | TMT-1 | 65.1 ± 12.3 | 49.6 ± 10.2 | 71.9 ± 10.1 | 95.9 ± 8.7 | 72.4 ± 9.4 | 43.3 ± 6.2 | 33.1 ± 5.8 | 22.0 ± 2.4 | 31.8 ± 3.8 |
|  | TMT-2 | 81.6 ± 22.8 | 81.0 ± 24.5 | 93.0 ± 12.1 | 89.5 ± 18.7 | 60.0 ± 8.1 | 39.1 ± 7.5 | 36.6 ± 8.0 | 35.0 ± 9.0 | 34.0 ± 4.4 |
| Female | Control | 54.9 ± 14.7 | 58.2 ± 13.6 | 63.4 ± 11.0 | 73.9 ± 10.0 | 54.8 ± 9.0 | 28.6 ± 2.2 | 29.9 ± 4.1 | 21.5 ± 3.1 | 26.8 ± 3.6 |
|  | TMT-1 | 65.1 ± 15.6 | 67.8 ± 14.1 | 57.2 ± 12.3 | 87.9 ± 13.1 | 51.8 ± 6.5 | 32.0 ± 7.2 | 29.2 ± 5.8 | 24.2 ± 2.7 | 30.4 ± 5.6 |
|  | TMT-2 | 67.9 ± 18.9 | 85.3 ± 21.0 | 81.8 ± 19.3 | 88.9 ± 28.3 | 59.0 ± 13.5 | 30.0 ± 7.7 | 32.4 ± 5.2 | 33.3 ± 5.7 | 27.6 ± 5.2 |
|  |  | Alcohol Intake (g/kg) |  |  |  |  |  |  |  |  |
| Male | Control | 0.2 ± 0.1 | 0.5 ± 0.1 | 1.3 ± 0.3 | 1.2 ± 0.2 | 1.2 ± 0.2 | 0.6 ± 0.1 | 1.1 ± 0.2 | 0.5 ± 0.1 | 0.6 ± 0.1 |
|  | TMT-1 | 0.2 ± 0.0 | 0.3 ± 0.1 | 0.9 ± 0.1 | 1.1 ± 0.1 | 1.3 ± 0.2 | 0.7 ± 0.1 | 0.8 ± 0.2 | 0.5 ± 0.1 | 0.7 ± 0.1 |
|  | TMT-2 | 0.2 ± 0.1 | 0.5 ± 0.2 | 1.2 ± 0.2 | 1.0 ± 0.2 | 1.0 ± 0.1 | 0.6 ± 0.1 | 0.8 ± 0.2 | 0.8 ± 0.2 | 0.7 ± 0.1 |
| Female | Control | 0.2 ± 0.0 | 0.5 ± 0.1 | 1.1 ± 0.2 | 1.1 ± 0.1 | 1.3 ± 0.2 | 0.8 ± 0.1 | 1.0 ± 0.1 | 0.7 ± 0.1 | 0.8 ± 0.1 |
|  | TMT-1 | 0.2 ± 0.1 | 0.6 ± 0.1 | 1.0 ± 0.2 | 1.5 ± 0.2 | 1.2 ± 0.2 | 0.7 ± 0.2 | 0.9 ± 0.2 | 0.7 ± 0.1 | 0.9 ± 0.1 |
|  | TMT-2 | 0.2 ± 0.1 | 0.7 ± 0.2 | 1.2 ± 0.3 | 1.3 ± 0.3 | 1.4 ± 0.4 | 0.7 ± 0.2 | 0.9 ± 0.1 | 0.9 ± 0.1 | 0.8 ± 0.1 |

Mean ± SEM; \* $p \leq 0.05$

### 3.1. TMT exposure produces distinct subgroups in male and female rats

Distribution plots (Figure 2A and B) represent the range of stress-reactive behaviors (digging and immobility) in male and female rats during TMT exposure. Males and females spent similar time engaged in defensive digging behavior during the TMT exposure, although the females had a greater range than the males (Figure 2A; Male: 0.0 to 140.6 sec; Female: 0 to 336.7 sec). Time spent immobile was significantly higher in males compared to females during the TMT exposure (Fig. 2B, t(32) =3.44, *p* < 0.05), indicating a sex difference in the engagement of immobility during the predator odor stress exposure. Using these stress-reactive behaviors, a digging/immobility ratio was calculated for each rat in the TMT group by dividing the total time spent digging by the total time spent immobile. As shown in Figure 1C, a median split of the scores was performed to divide the range of ratio scores into two TMT-subgroups for each sex; ratio scores below the median were classified as TMT-subgroup 1 and ratio scores above the median were classified as TMT-subgroup 2 in both male and female rats. From this point forward, males and females were analyzed separately.

**Figure 2.**
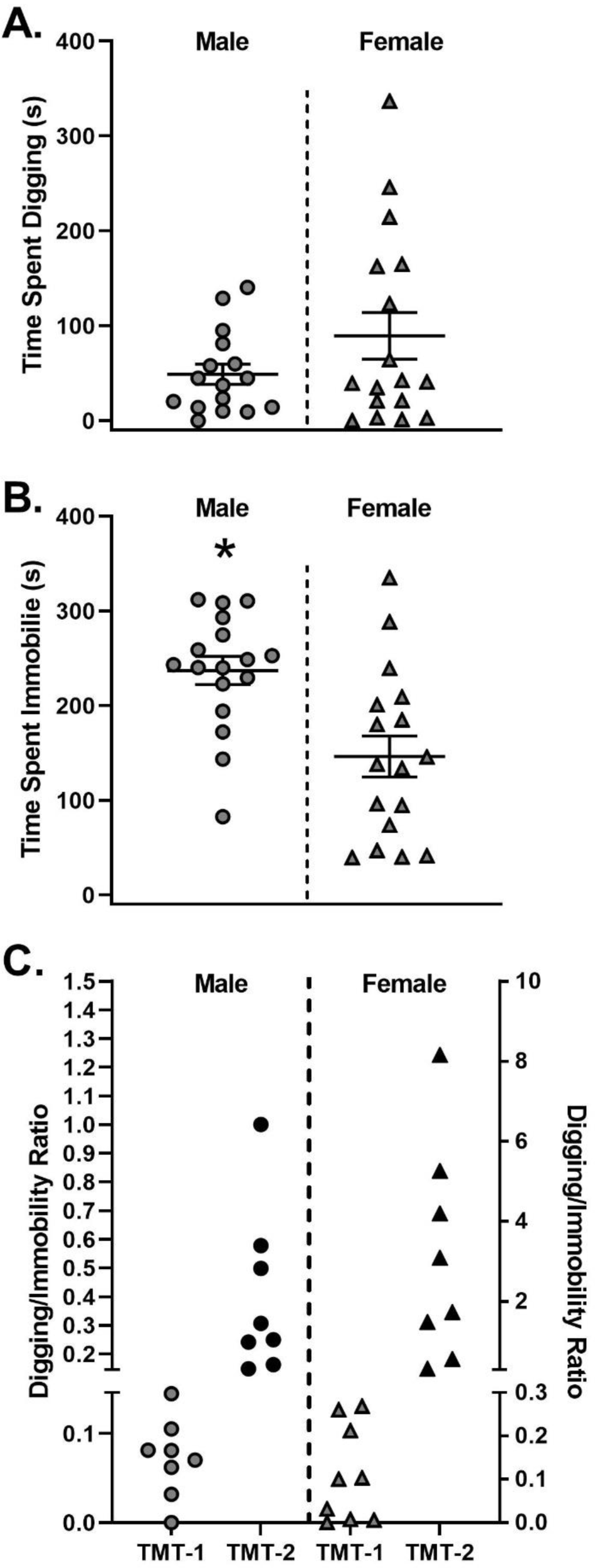
Distribution plots to illustrate range of stress-reactive behaviors between male and female rats exposed to TMT. (A) Male and female rats exposed to TMT showed no difference in total defensive digging. (B) Male rats exposed to TMT spend significantly more total time immobile compared to female rats exposed to TMT. (C) Representation of digging/immobility scores in male and female TMT-groups 1 and 2 with median split. In both male and female rats, TMT-subgroup 1 had lower digging/immobility scores compared to TMT-subgroup 2, \**p* < 0.05.

### 3.2. TMT-subgroups in male and female rats display distinct behavioral phenotypes during TMT exposure

During the TMT exposure, there was a significant group difference in total time spent digging in males and females (Fig. 3A, Males: F (2,23) = 27.92, *p* < 0.05; Fig. 3H, Females: F (2,26) = 23.50, *p* < 0.05). Males and females in TMT-subgroup 2 engaged in significantly more digging compared to TMT-subgroup 1 and controls (*p* < 0.05). Cumulative digging was used to examine the pattern of digging across time. The two-way RM ANOVA showed a significant main effect of group (Fig. 3B, Males: F (2,23) = 27.42, *p* < 0.05; Fig. 3I, Females: F (2,26) = 23.41, *p* < 0.05), time (Fig. 3B, Males: F (9,207) = 50.08, *p* < 0.05; Fig. 3I, Females: F (9,234) = 37.86, *p* < 0.05), and a significant group by time interaction (Fig. 3B, Males: F (18,207) = 25.57, *p* < 0.05; Fig. 3I, Females: F (18,234) = 22.21, *p* < 0.05). In males and females, digging in the TMT-subgroup 2 was significantly greater than controls and TMT 1 by minute 3 and throughout the remainder of the exposure (*p* < 0.05), indicating that the emergence of digging behavior early in the session.

**Figure 3.**
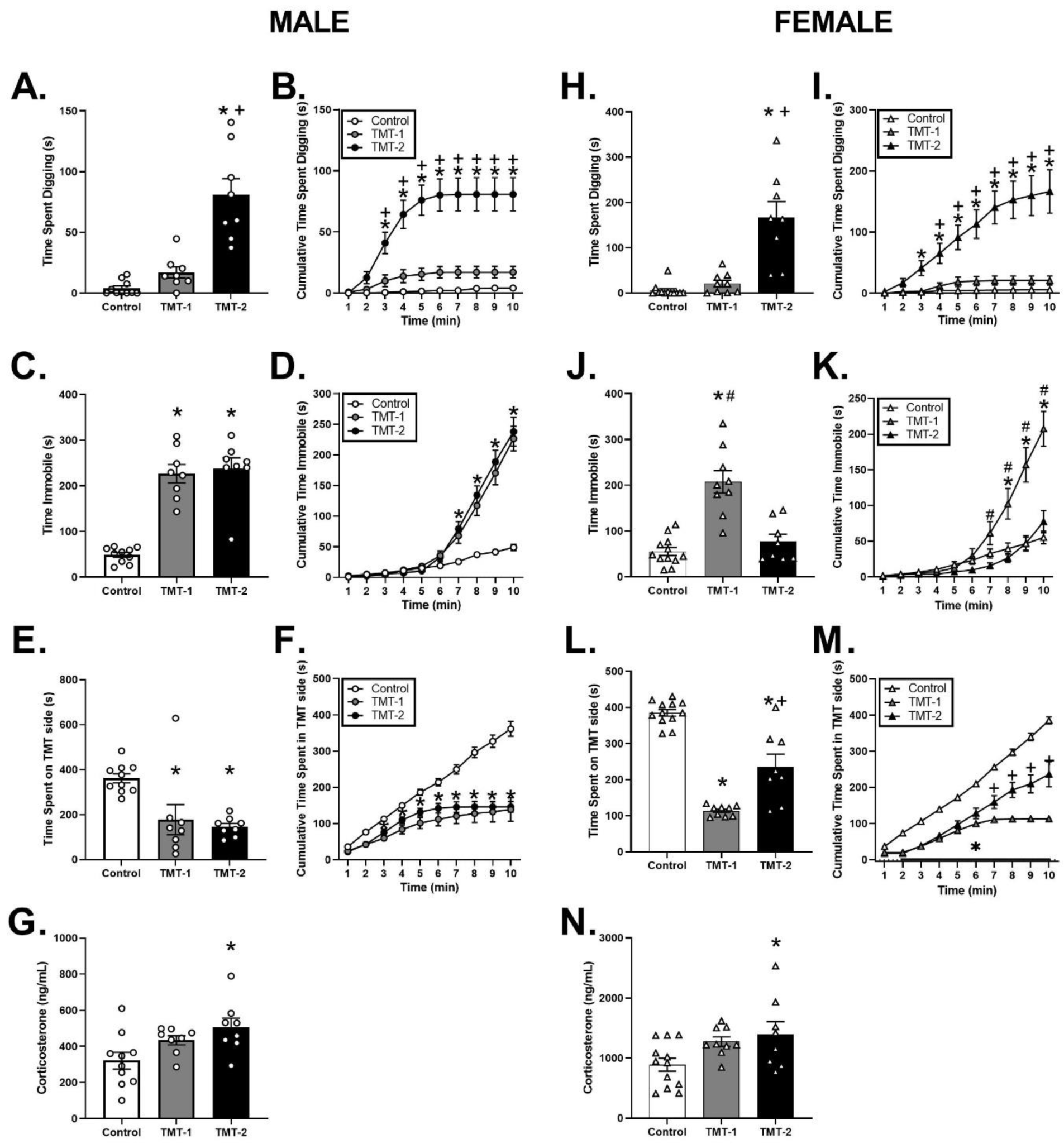
Effects of TMT exposure on stress-reactive behaviors in male and female TMT-subgroups. Male rats in TMT-subgroup 2 performed significantly more defensive digging compared to TMT-subgroup 1 and controls (A,) as well as across 10 min of the TMT exposure (B). Male rats in TMT-subgroup 1 and 2 exhibited significantly more time immobile (C) and time spent on TMT side (E) compared to controls. Across 10 min of the TMT exposure, males in TMT-subgroup 1 and 2 spent significantly more time immobile (D) and decreased time spent on TMT side (F) compared to controls. Male rats in TMT-subgroup 2 exhibited higher plasma corticosterone compared to controls (G). Female rats in TMT-subgroup 2 performed significantly more defensive digging compared to controls and TMT-subgroup 1 (H), as well as across the 10 min TMT exposure (I). Female rats in TMT subgroup 1 exhibited significantly more time immobile compared to controls and TMT-subgroup 2 (J), as well as across 10 min of the TMT exposure (K). Female rats in both TMT-subgroups 1 and 2 spent significantly less time on TMT side compared to controls (L), as well as across the 10 min TMT exposure (M). Female rats in TMT-subgroup 2 exhibited higher plasma corticosterone levels compared to controls (N). * p < 0.05 vs. controls, # p < 0.05 vs. TMT-2, + p < 0.05 vs. TMT-1

As shown in Figure 3C and J, there was a significant main effect of TMT on total time immobile in males (F (2,23) = 42.32, *p* < 0.05) and females (F(2,26) = 25.17, *p* < 0.05). In males, TMT-subgroup 1 and 2 spent greater time immobile compared to controls (*p* < 0.05). However, in females, TMT-subgroup 1 spent greater time immobile compared to TMT-subgroup 2 and controls (*p* < 0.05). To examine the pattern of immobility over time, a two-way RM ANOVA of showed a significant main effect of group (Fig. 3D, Males: F (2,23) = 24.70, *p* < 0.05; Fig. 3K, Females: F (2,26) = 10.51, *p* < 0.05), time (Fig. 3D, Males: F (9,207) = 215.2, *p* < 0.05; Fig. 3K, Females: F (9,234) = 90.04, *p* < 0.05), and a significant time by group interaction (Fig. 3D, Males: F (18,207) = 32.01, *p* < 0.05; Fig. 3K, Females: F (18,234) = 19.31, *p* < 0.05). In males, time immobile in both TMT-subgroups was significantly greater than controls by minute 7 throughout the remainder of the exposure (*p* < 0.05). In females, time immobile in TMT-subgroup 1 was significantly greater than controls and TMT-subgroup 2 by minute 7 and controls by minute 8 throughout the remainder of the exposure (*p* < 0.05), indicating that immobility behavior emerged later in the session.

Time spent on the TMT side of the chamber was examined as a measure of avoidance behavior. There was a significant main effect of TMT in both males (Fig. 3E, F (2,23) = 9.43, *p* < 0.05) and females (Fig. 3L, F (2,26) = 61.81, *p* < 0.05). In males, TMT-subgroups 1 and 2 spent significantly less time on the TMT side of the chamber compared to controls (*p* < 0.05). In females, TMT-subgroup 1 spent significantly less time on the TMT side of the chamber compared to TMT-subgroup 2 and controls, while TMT-subgroup 2 avoided the TMT side of the chamber less than controls only (*p* < 0.05). Analysis of the distribution of time on the TMT side showed a significant main effect of group (Fig. 3F, Males: F (2,23) = 24.54, *p* < 0.05; Fig. 3M, Females: F (2,26) = 89.27, *p* < 0.05) and time (Fig. 3F, Males: F (9,207) = 172.80, *p* < 0.05; Fig. 3M, Females: F (9,234) = 364.4, *p* < 0.05), as well as a significant time by group interaction (Fig. 3F, Males: F (18,207) = 24.93, *p* < 0.05; Fig. 3M, Females: F (18,234) = 35.76, *p* < 0.05). In males, time spent on TMT side in TMT-subgroup 1 was significantly lower by minute 3 compared to controls and in TMT-subgroup 2 lower by minute 5 compared to controls throughout the remainder of the session (*p* < 0.05). In females, time spent on TMT side in TMT subgroups 1 and 2 was significantly lower from minute 2 and throughout the remainder of the session (*p* < 0.05), while TMT-subgroup 2 spent significantly more time on TMT side compared to TMT-subgroup 1 from minute 7 throughout the remainder of the session, indicating that rats gradually increased avoidance of the side of the chamber the TMT was located throughout the exposure.

Fecal boli production during TMT exposure and corticosterone levels following TMT exposure were used as indices of physiological responses to TMT exposure. Fecal boli did not differ in males (F (2,23) = 0.89, *p* > 0.05; Control: 1.70 ± 0.90, TMT-1: 3.13 ± 0.99, TMT-2: 3.25 ± 0.94). However, in females, there was a significant main effect of TMT (F (2,26) = 7.03, *p* < 0.05), data not shown) with both TMT-subgroups 1 and 2 showing more fecal boli compared to controls (*p* < 0.05). In males and females, plasma corticosterone levels were significantly increased following TMT exposure (Fig. 3G, Males: F (2,23) = 4.73, *p* < 0.05; Fig. 3N, Females: F (2,26) = 4.12, *p* < 0.05). In both males and females, TMT-subgroup 2 showed significantly higher plasma corticosterone levels compared to controls (*p* < 0.05). Overall, these data show specific behavioral phenotypes in male and female TMT-subgroups in during the TMT predator odor exposure.

### 3.3. TMT-subgroups in male and female rats display distinct behavioral phenotypes during context re-exposure

Analysis of digging during context re-exposure showed a significant main effect of TMT in males (Fig. 4A, F (2,23) = 5.81, *p* < 0.05) and females (Fig. 4H, F (2,26) = 10.84, *p* < 0.05). Male and female rats in TMT-subgroup 2 engaged in significantly more defensive digging compared to controls (*p* < 0.05), and female TMT-subgroup 2 engaged in significantly more defensive digging compared TNT-subgroup 1. In contrast, no difference in digging behavior was observed between TMT subgroup 1 and controls in either males or females (*p* > 0.05). Examination of cumulative digging across time showed a significant main effect of group (Fig. 4B, Males: F (2,23) = 5.17, *p* < 0.05; Fig. 5I, Females: F (2,26) = 11.20, *p* < 0.05), time (Males: F (9,207) = 14.56, *p* < 0.05; Females: F (9,234) = 18.06, *p* < 0.05), and a significant group by time interaction (Males: F (18,207) = 5.63, *p* < 0.05; Females: F (18,234) = 7.26, *p* < 0.05). Male TMT-subgroup 2 engaged in significantly more defensive digging compared to controls from minute 5 and TMT-subgroup 1 from minute 8 through the duration of the exposure (*p*<0.05). In females, TMT-subgroup 2 engaged in significantly more defensive digging compared to controls and TMT-subgroup 1 from minute 5 through the duration of the exposure (*p*<0.05), indicating a similar pattern as digging during TMT exposure.

**Figure 4.**
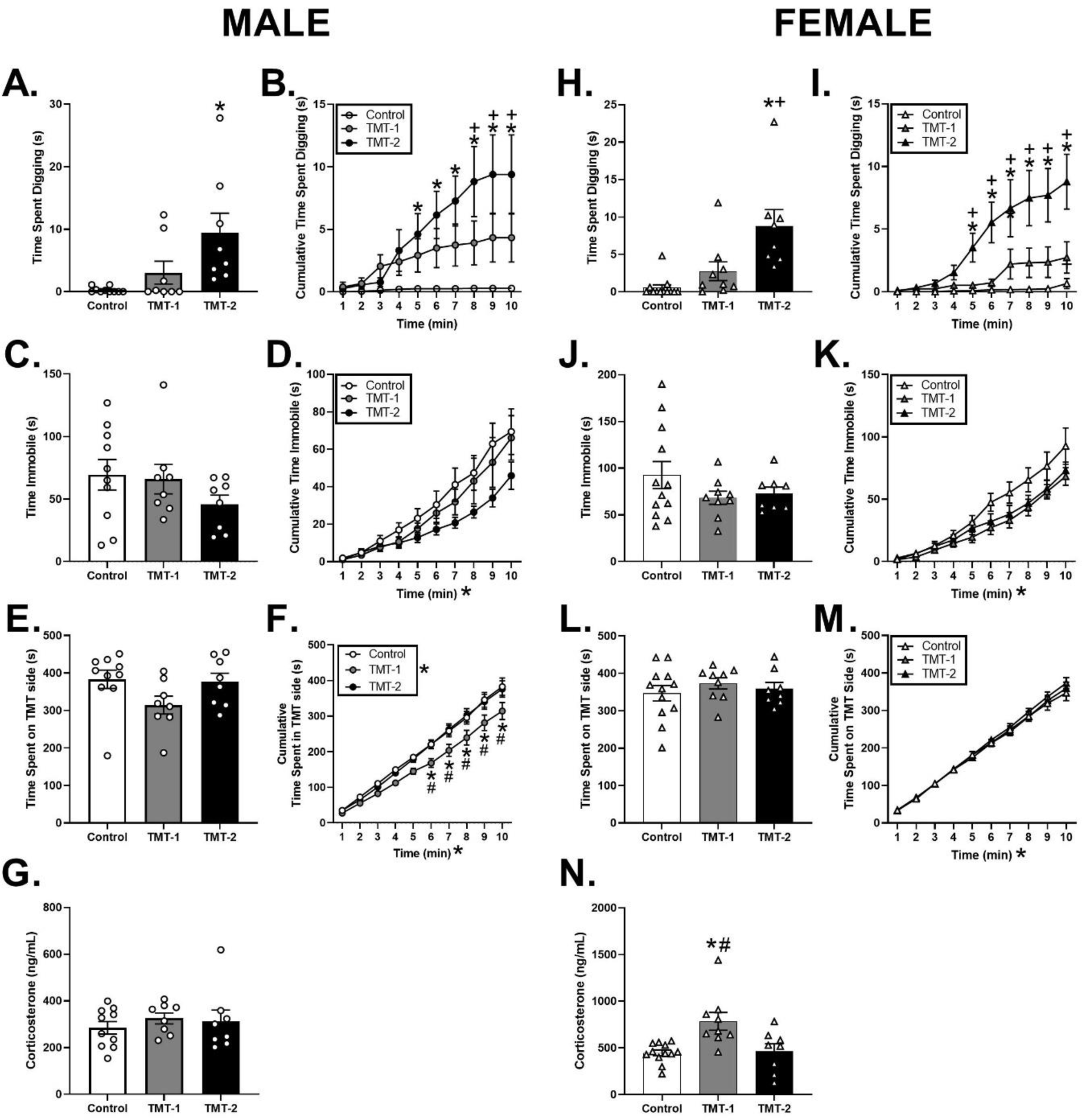
Reactivity to predator odor-paired context in male and female TMT-subgroups. During context re-exposure, males and females performed significant more digging. Specifically, male (A) and female (H) TMT-subgroup 2 performed greater total time digging, as well as across time (B, Male; I, Female) compared to controls. Males and females previously exposed to TMT showed no changes in total time immobile (Males, C; Females, J) or across time (Males, D; Females, K) compared to controls. There were differences between males TMT-subgroups and controls for total time spent on the TMT side; however, across time, male TMT-subgroup 1 spent significantly less time on the TMT side compared to controls and TMT-subgroup 2 (E). There were no differences in time spent on TMT side in females (M). There were no differences in plasma corticosterone after context re-exposure in males (G); however, in females, TMT-subgroup 1 exhibited greater plasma corticosterone levels compared to controls and TMT-subgroup 2. * p < 0.05 vs. controls, # p < 0.05 vs. TMT-2, + p < 0.05 vs. TMT-1

**Figure 5.**
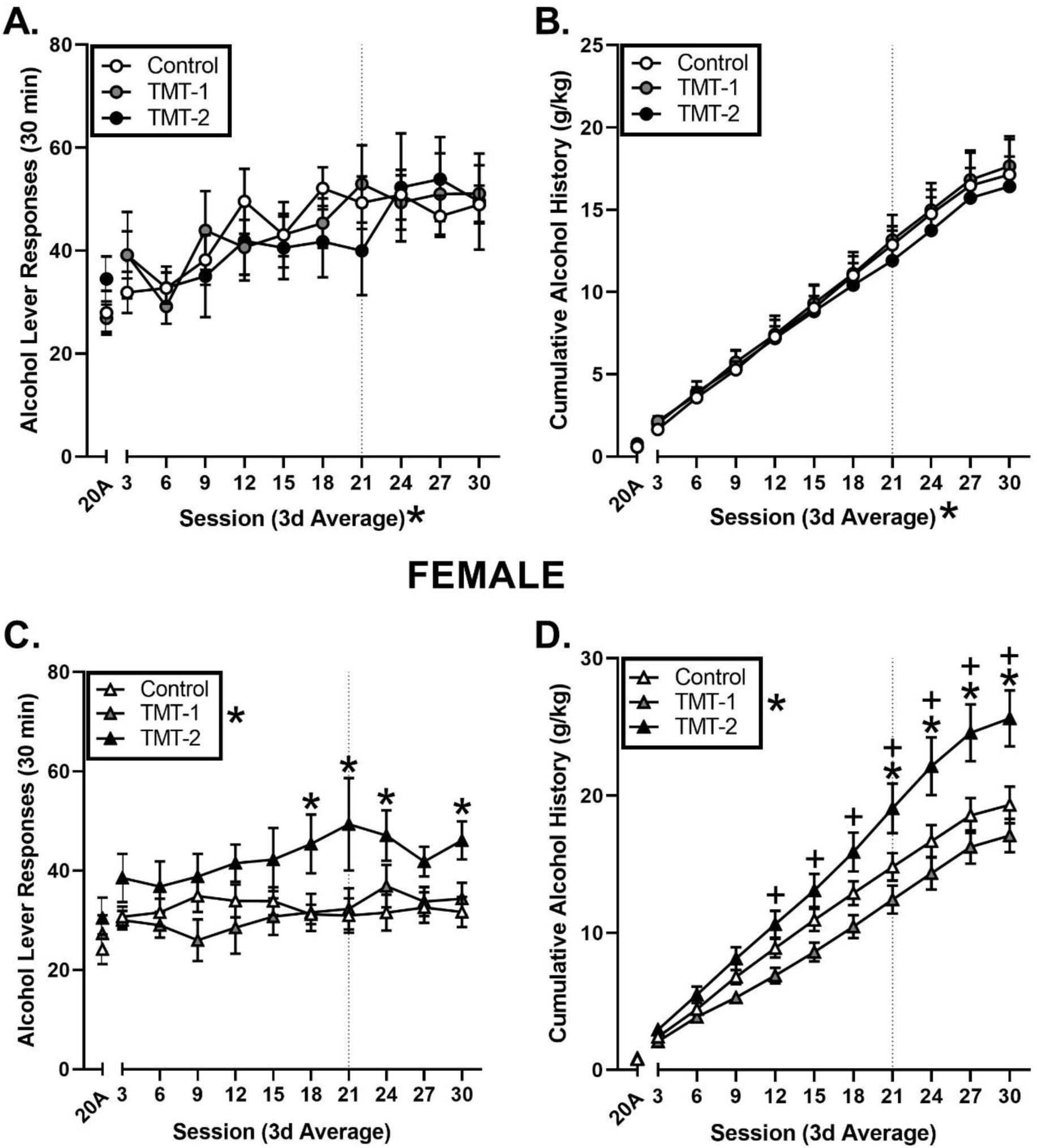
Maintenance of alcohol self-administration following TMT exposure in male and female TMT-subgroups. Males: TMT-subgroups 1 and 2 exhibited no significant increases in alcohol lever responses (A) or cumulative alcohol history (B). Females: TMT-subgroup 2 performed significantly greater alcohol lever responses compared to controls (C) and cumulative alcohol intake (D) compared to TMT-subgroup 1 and controls. * p < 0.05 vs. controls, # p < 0.05 vs. TMT-2, + p < 0.05 vs. TMT-1. Dotted line denotes moving to MWF sessions.

Examination of total time immobile during context re-exposure showed no significant main effect of TMT in males (Fig. 4C) and females (Fig. 4J) (*p* > 0.05). Analysis of cumulative time immobile across time showed a significant main effect of time in male (Fig. 4D, F (9,207) = 59.52, *p* < 0.05) and female (Fig. 4K, F (9,234) = 95.89, *p* < 0.05), but no significant main effect of group or significant group by time interaction (*p* > 0.05), indicating immobility was not a behavior indicative of context reactivity.

Analysis of time spent on the side in which TMT was located showed no significant main effect of TMT in males (Fig. 4E, *p* > 0.05) or females (Fig. 4L, *p* < 0.05). However, two-way RM ANOVA of cumulative time spent on TMT side across time in males showed a significant group x time interaction (Fig. 4F, F (18,207) = 1.76, *p* < 0.05), a significant main effect of group (F (2,23) = 4.49, *p* < 0.05) and time (F (9,207) = 423.7, *p* < 0.05). Male TMT-subgroup 1 spent significantly less time on the side in which TMT was located compared to controls and TMT-subgroup 2 from minute 6 throughout the duration of the exposure. In contrast, across time analysis in females showed no main effect of group, time or significant interaction (*p* > 0.05), indicating a subset of males avoided the TMT side during context re-exposure compared to females.

Examination of fecal boli production during context re-exposure showed a significant main effect of TMT group in males (Fig. 4G, F (2,23) = 4.23, *p* < 0.05, Controls: 0.00 ± 0.00, TMT-1: 1.25 ± 0.65, TMT-2: 0.00 ± 0.00), but not in females (*p*> 0.05, Controls: 0.00 ± 0.00, TMT-1: 0.77 ± 0.47, TMT-2: 0.00 ± 0.00), with more fecal boli in TMT-subgroup 1 compared to controls in males (*p* < 0.05).

In males, there was no group difference in plasma corticosterone levels (Fig. 4G, *p* > 0.05), but females showed a significant difference (Fig. 4N, F (2,26) = 7.96, *p* < 0.05), with higher levels in TMT-subgroup 1 compared to controls and TMT-subgroup 2 (*p* < 0.05).

### 3.4. TMT increases hyperarousal behavior in a subset of female but not male rats

Analysis of anxiety-like behavior during the light dark test and the zero maze showed no significant differences between TMT-subgroups and controls in male and female rats (Table 2). However, in the acoustic startle test, there was a significant main effect of TMT group in females (Table 4; F (2,26) = 6.68, *p* < 0.05), in which female rats in TMT-subgroup 1 exhibited a higher average startle amplitude compared to controls and TMT-subgroup 2 (*p*<0.05), indicating females show greater hyperarousal behavior after TMT exposure.

**Table 2.**
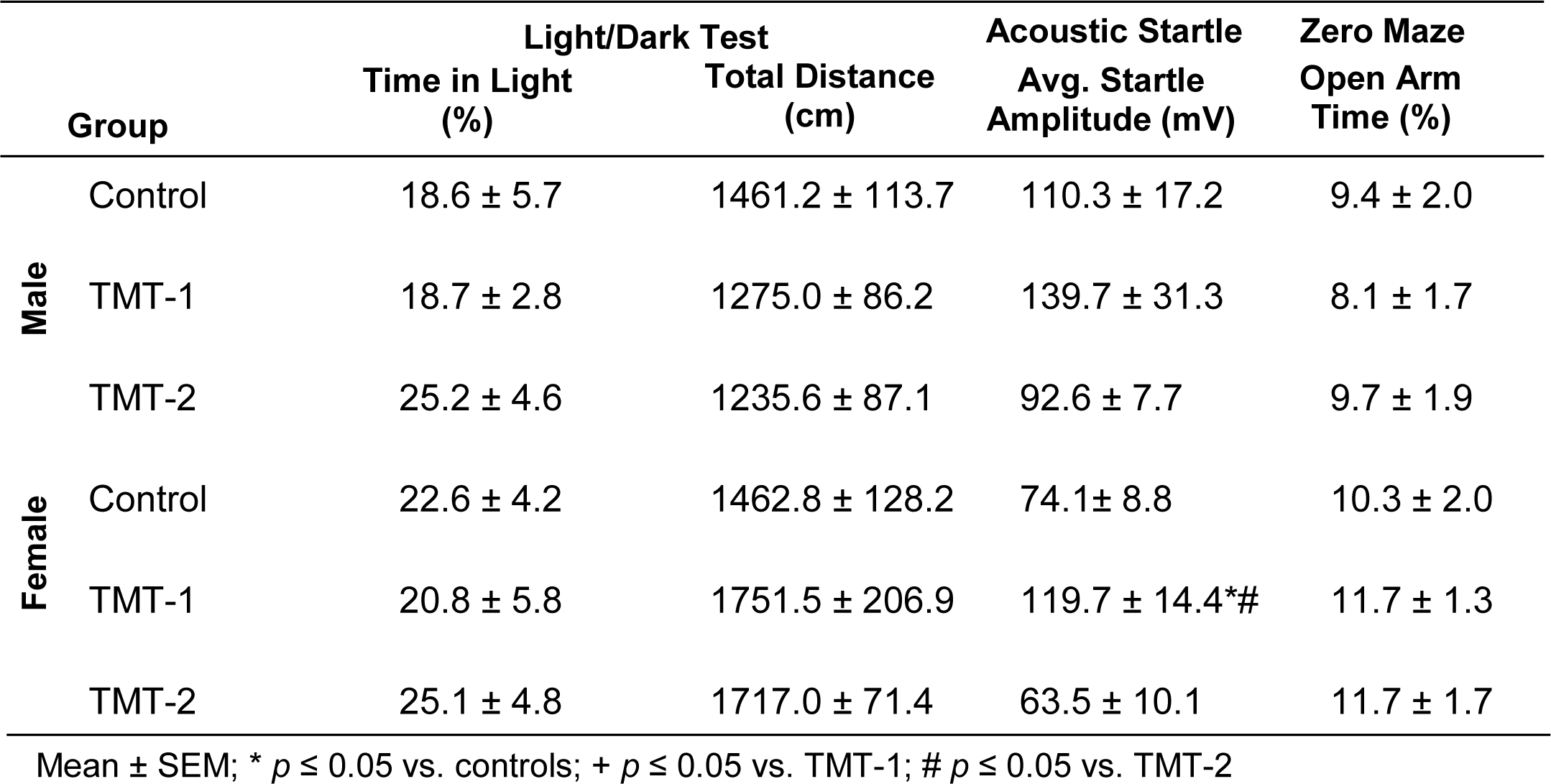
Behavioral Screen Data

### 3.5. TMT increases alcohol self-administration in female but not male rats

During alcohol self-administration post-TMT exposure, male rats showed a significant main effect of session for alcohol lever responses (Fig. 5A, F (10,230) = 11.13, *p* < 0.05) and cumulative alcohol drinking history (Fig. 5B, F (10,230) = 219.5, *p* < 0.05), indicating increased drinking over time. However, there was no main effect of group (lever responses or alcohol intake, *p* > 0.05) or significant interaction (lever responses or alcohol intake, *p* > 0.05). Session averages for alcohol intake (g/kg) are shown in Table 3). These results show that TMT exposure did not affect ongoing alcohol self-administration in male rats following TMT exposure.

**Table 3.**
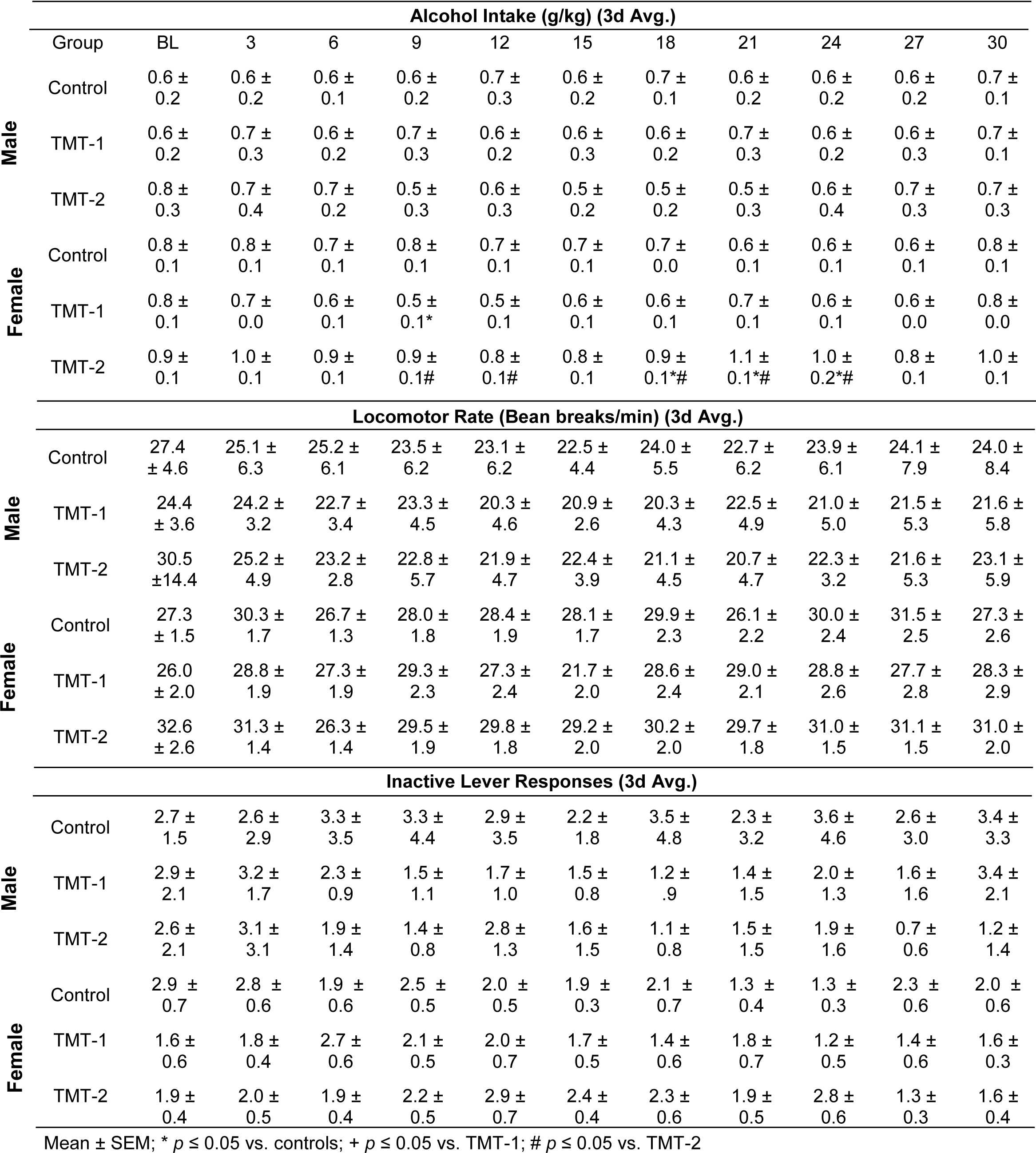
Alcohol Intake (g/kg), Locomotor rate, and Inactive Lever Responses during 30 sessions of self-administration

|  |  | Alcohol Intake (g/kg) (3d Avg.) |  |  |  |  |  |  |  |  |  |  |
| --- | --- | --- | --- | --- | --- | --- | --- | --- | --- | --- | --- | --- |
| Male | Group | BL | 3 | 6 | 9 | 12 | 15 | 18 | 21 | 24 | 27 | 30 |
|  | Control | 0.6 ± 0.2 | 0.6 ± 0.2 | 0.6 ± 0.1 | 0.6 ± 0.2 | 0.7 ± 0.3 | 0.6 ± 0.2 | 0.7 ± 0.1 | 0.6 ± 0.2 | 0.6 ± 0.2 | 0.6 ± 0.2 | 0.7 ± 0.1 |
|  | TMT-1 | 0.6 ± 0.2 | 0.7 ± 0.3 | 0.6 ± 0.2 | 0.7 ± 0.3 | 0.6 ± 0.2 | 0.6 ± 0.3 | 0.6 ± 0.2 | 0.7 ± 0.3 | 0.6 ± 0.2 | 0.6 ± 0.3 | 0.7 ± 0.1 |
|  | TMT-2 | 0.8 ± 0.3 | 0.7 ± 0.4 | 0.7 ± 0.2 | 0.5 ± 0.3 | 0.6 ± 0.3 | 0.5 ± 0.2 | 0.5 ± 0.2 | 0.5 ± 0.3 | 0.6 ± 0.4 | 0.7 ± 0.3 | 0.7 ± 0.3 |
| Female | Control | 0.8 ± 0.1 | 0.8 ± 0.1 | 0.7 ± 0.1 | 0.8 ± 0.1 | 0.7 ± 0.1 | 0.7 ± 0.1 | 0.7 ± 0.0 | 0.6 ± 0.1 | 0.6 ± 0.1 | 0.6 ± 0.1 | 0.8 ± 0.1 |
|  | TMT-1 | 0.8 ± 0.1 | 0.7 ± 0.0 | 0.6 ± 0.1 | 0.5 ± 0.1* | 0.5 ± 0.1 | 0.6 ± 0.1 | 0.6 ± 0.1 | 0.7 ± 0.1 | 0.6 ± 0.1 | 0.6 ± 0.0 | 0.8 ± 0.0 |
|  | TMT-2 | 0.9 ± 0.1 | 1.0 ± 0.1 | 0.9 ± 0.1 | 0.9 ± 0.1# | 0.8 ± 0.1# | 0.8 ± 0.1 | 0.9 ± 0.1*# | 1.1 ± 0.1*# | 1.0 ± 0.2*# | 0.8 ± 0.1 | 1.0 ± 0.1 |
|  |  | Locomotor Rate (Bean breaks/min) (3d Avg.) |  |  |  |  |  |  |  |  |  |  |
| Male | Control | 27.4 ± 4.6 | 25.1 ± 6.3 | 25.2 ± 6.1 | 23.5 ± 6.2 | 23.1 ± 6.2 | 22.5 ± 4.4 | 24.0 ± 5.5 | 22.7 ± 6.2 | 23.9 ± 6.1 | 24.1 ± 7.9 | 24.0 ± 8.4 |
|  | TMT-1 | 24.4 ± 3.6 | 24.2 ± 3.2 | 22.7 ± 3.4 | 23.3 ± 4.5 | 20.3 ± 4.6 | 20.9 ± 2.6 | 20.3 ± 4.3 | 22.5 ± 4.9 | 21.0 ± 5.0 | 21.5 ± 5.3 | 21.6 ± 5.8 |
|  | TMT-2 | 30.5 ± 14.4 | 25.2 ± 4.9 | 23.2 ± 2.8 | 22.8 ± 5.7 | 21.9 ± 4.7 | 22.4 ± 3.9 | 21.1 ± 4.5 | 20.7 ± 4.7 | 22.3 ± 3.2 | 21.6 ± 5.3 | 23.1 ± 5.9 |
| Female | Control | 27.3 ± 1.5 | 30.3 ± 1.7 | 26.7 ± 1.3 | 28.0 ± 1.8 | 28.4 ± 1.9 | 28.1 ± 1.7 | 29.9 ± 2.3 | 26.1 ± 2.2 | 30.0 ± 2.4 | 31.5 ± 2.5 | 27.3 ± 2.6 |
|  | TMT-1 | 26.0 ± 2.0 | 28.8 ± 1.9 | 27.3 ± 1.9 | 29.3 ± 2.3 | 27.3 ± 2.4 | 21.7 ± 2.0 | 28.6 ± 2.4 | 29.0 ± 2.1 | 28.8 ± 2.6 | 27.7 ± 2.8 | 28.3 ± 2.9 |
|  | TMT-2 | 32.6 ± 2.6 | 31.3 ± 1.4 | 26.3 ± 1.4 | 29.5 ± 1.9 | 29.8 ± 1.8 | 29.2 ± 2.0 | 30.2 ± 2.0 | 29.7 ± 1.8 | 31.0 ± 1.5 | 31.1 ± 1.5 | 31.0 ± 2.0 |
|  |  | Inactive Lever Responses (3d Avg.) |  |  |  |  |  |  |  |  |  |  |
| Male | Control | 2.7 ± 1.5 | 2.6 ± 2.9 | 3.3 ± 3.5 | 3.3 ± 4.4 | 2.9 ± 3.5 | 2.2 ± 1.8 | 3.5 ± 4.8 | 2.3 ± 3.2 | 3.6 ± 4.6 | 2.6 ± 3.0 | 3.4 ± 3.3 |
|  | TMT-1 | 2.9 ± 2.1 | 3.2 ± 1.7 | 2.3 ± 0.9 | 1.5 ± 1.1 | 1.7 ± 1.0 | 1.5 ± 0.8 | 1.2 ± .9 | 1.4 ± 1.5 | 2.0 ± 1.3 | 1.6 ± 1.6 | 3.4 ± 2.1 |
|  | TMT-2 | 2.6 ± 2.1 | 3.1 ± 3.1 | 1.9 ± 1.4 | 1.4 ± 0.8 | 2.8 ± 1.3 | 1.6 ± 1.5 | 1.1 ± 0.8 | 1.5 ± 1.5 | 1.9 ± 1.6 | 0.7 ± 0.6 | 1.2 ± 1.4 |
| Female | Control | 2.9 ± 0.7 | 2.8 ± 0.6 | 1.9 ± 0.6 | 2.5 ± 0.5 | 2.0 ± 0.5 | 1.9 ± 0.3 | 2.1 ± 0.7 | 1.3 ± 0.4 | 1.3 ± 0.3 | 2.3 ± 0.6 | 2.0 ± 0.6 |
|  | TMT-1 | 1.6 ± 0.6 | 1.8 ± 0.4 | 2.7 ± 0.6 | 2.1 ± 0.5 | 2.0 ± 0.7 | 1.7 ± 0.5 | 1.4 ± 0.6 | 1.8 ± 0.7 | 1.2 ± 0.5 | 1.4 ± 0.6 | 1.6 ± 0.3 |
|  | TMT-2 | 1.9 ± 0.4 | 2.0 ± 0.5 | 1.9 ± 0.4 | 2.2 ± 0.5 | 2.9 ± 0.7 | 2.4 ± 0.4 | 2.3 ± 0.6 | 1.9 ± 0.5 | 2.8 ± 0.6 | 1.3 ± 0.3 | 1.6 ± 0.4 |

Mean ± SEM; \* $p \leq 0.05$ vs. controls; + $p \leq 0.05$ vs. TMT-1; # $p \leq 0.05$ vs. TMT-2

In contrast, in female rats there was a significant main effect of group on alcohol lever responses (Fig. 5C, F (2,26) = 4.69, *p* < 0.05), in which TMT-subgroup 2 had significantly greater alcohol lever responses compared to TMT-subgroup 1 and controls. There was no main effect of session and no interaction. However, to further explore the group difference in self-administration and based on the a priori hypothesis that TMT-subgroup 2 would show increases in alcohol self-administration, planned comparisons tests using Bonferroni correction for multiple comparisons were conducted across each alcohol session. TMT-subgroup 2 had higher alcohol lever responses compared to controls from session 18 through the duration of alcohol self-administration (*p* < 0.05). Furthermore, examination of cumulative alcohol drinking history, showed a significant main effect of group (Fig. 5D, F (2,26) = 6.56, *p* < 0.05), in which TMT-subgroup 2 had significantly greater alcohol intake compared to TMT-subgroup 1 and controls. There was also a significant main effect of session (F (9,234) = 415.10, *p* < 0.05), as well as a significant session by group interaction (F (18,234) = 6.71, *p* < 0.05). Post hoc comparisons showed that TMT-subgroup 2 had higher alcohol intake compared to controls from session 21 through duration of alcohol self-administration, as well as higher alcohol intake compared to TMT-subgroup 1 from session 12 through the duration of alcohol self-administration (*p* ≤ 0.05). Session averages for alcohol intake (g/kg) are shown in Table 3). There no changes in locomotor behavior or inactive lever responses between controls and TMT-subgroups in male and female rats (Table 3). These results indicate TMT subgroup classifications using the digging immobility ratio during TMT predicts increases in alcohol self-administration in female, but not male rats.

## 4. Discussion

In the current study we sought to examine if individual differences in stress-reactive behaviors during TMT exposure (a ratio of defensive digging/immobility) could predict subsequent increases in context reactivity, hyperarousal, anxiety-like behavior and alcohol self-administration in male and female rats. The results demonstrate several important findings. First, during TMT exposure, male and female Long-Evans rats engage in defensive digging and immobility behaviors, but males spend more time immobile than females, and females show a larger range in defensive digging behavior (though overall time spent digging was not different from males). Second, male and female rats, with higher ratio scores of digging/immobility (TMT-subgroup 2) showed increased CORT levels during TMT exposure (elevated stress response) compared to controls, indicating an enhanced HPA-axis stress response associated with high digging/immobility ratio scores. Third, male and female rats in TMT-subgroup 2 showed enhanced reactivity to the TMT-paired context (i.e., increased defensive digging), suggesting a contextually-cued stress memory in rats that showed higher stress responsivity during TMT exposure. Interestingly, a subset of females who engaged in passive coping behavior (i.e., immobility; TMT-subgroup 1) during TMT exhibited higher corticosterone levels during context re-exposure, suggestive of a contextual-induced physiological stress response. Fourth, female rats with lower digging/immobility ratio scores (TMT-subgroup 1) showed hyperarousal as measured by acoustic startle, but this was not observed in males. Lastly, female but not male rats, in TMT-subgroup 2 (high digging/low immobility ratio) showed escalations in alcohol self-administration that emerged 24 days (12 sessions) following TMT exposure, indicating a lasting consequence of stressor exposure. Together, these data suggest that stress-reactive behaviors during predator odor stressor exposure using TMT may be helpful in understanding individual resilience/susceptibility to stress and understanding the impact of stress on escalations in alcohol drinking.

TMT elicits innate fear and stress-related behavioral responses including freezing [34, 40], immobility [27]; [36] and avoidance behavior [27]; [35, 36, 41]. Here, bedding was present in the TMT exposure context so that we could examine an additional stress-reactivity behavior, defensive digging [27]. Defensive digging during TMT exposure involved the rat actively moving bedding material toward the corner of the chamber directly below the basket that held the TMT. Previous studies have defined such behavior as defensive burying [29, 30]; however, as the rat was not actively burying an object but burrowing the head, forepaws or entire body into the bedding and performing shoveling movements, we categorized the behavior as defensive digging. Here, we show male and female Long-Evans rats engaged in similar amount of total time digging during the TMT exposure (Figure 2B), which is consistent with previous work showing that male and female rats respond to TMT with similar increases in frequency and duration of defensive burying [42] and defensive digging [35]. Although overall time spent digging was not different from males, there was a larger range of individual variability in females, suggestive of heterogeneous proactive coping behavior to TMT. Interestingly, during TMT exposure, both sexes showed a shift in behavioral responses, transitioning from defensive digging to immobility and avoidance behavior, with male rats showing greater total immobility than females. This transition has been described as a stress-induced shift in coping behavior, in which rats will shift from active to passive coping strategies [31, 43].

Males engaging in greater immobility behavior than females is in accordance with previous studies showing freezing behavior is a male-typical behavior that has been frequently studied in fear conditioning and extinction studies [44, 45]. Male rats exhibit greater conditioned freezing, as well as immobility during single forced swim tests compared to females [46]. Studies using TMT show male rats engage in diverse defensive behaviors including immobility and avoidance when exposed in larger chambers, whereas in a smaller inescapable, rats engage in freezing behavior, suggestive of a fear response [34]. In one study, male rats exposed to TMT engaged in similar increases in immobility and decreased preference for the side containing TMT (i.e. avoidance) compared to females [35]. Here, we show male and female rats engage in similar levels of defensive digging behavior; however, males engage in greater immobility behavior throughout the duration of the TMT exposure compared to females.

To focus on individual differences in stress-reactivity, rats were divided into stress-reactivity subgroups based on a ratio of their digging and immobility behaviors during TMT exposure using a median split. This produced distinct subgroups within TMT exposed male and female rats: TMT-subgroup 1 (low digging/ immobility ratio), TMT-subgroup 2 (high digging/ immobility ratio). We specifically chose digging and immobility behavior as an index of stress-reactivity because they are both innate stress responsive behaviors but represent two distinctly different behavioral coping strategies [29, 37, 38].

In male rats, TMT-subgroup 1 engaged in little to no digging behavior but had high levels of immobility compared to TMT-subgroup 2 that showed high levels of both digging and immobility. The latter was an unexpected finding because rats that engage in more time digging typically show less time immobile due to the overall increase in activity. Furthermore, male TMT-subgroup 2 showed increased plasma corticosterone levels. Interestingly, females in TMT-subgroup 1 engaged in low levels of digging and high levels of immobility, while TMT-subgroup 2 engaged in high levels of digging and low levels of immobility, and had increased plasma corticosterone levels. We interpret differences in female stress-reactive behaviors between the TMT-subgroups as representing two distinct forms of coping responses, specifically with TMT-subgroup 1 engaging in more passive responses (i.e., immobility) and TMT-subgroup 2 engaging in active coping responses (i.e., digging). Male rats, regardless of TMT-subgroups, engaged in high levels of immobility behavior indicative of passive coping, which is a typical behavioral phenotype exhibited in males when responding to stress [44]. While we were able to capture two TMT-subgroups in male rats using the digging/immobility ratio score, it is clear that these ratio scores for both male TMT-subgroups represent a higher proportion of immobility behavior compared to digging behavior. Furthermore, increased plasma corticosterone was observed in both male and females in TMT-subgroup 2. Given that increased digging was a similar behavioral response in this subgroup across sex, this suggests that engagement in digging behavior may be a predictor of corticosterone response to TMT.

Two out of the four clusters of diagnostic criteria for PTSD within the DSM-5 include 1) intrusive distressing memories of the traumatic event(s) and 2) persistent avoidance of the stimuli associated with the traumatic event(s). Understanding the behavioral and neurobiological mechanisms associated with these persistent and long-lasting symptoms can be invaluable tools for effective treatment and prevention strategies for PTSD. In animal models, re-exposing animals to stress-related stimuli or the environment in which the stressor was presented can induce contextual fear or stress responses that can serve as an index of memory of that context. Male and female rats in TMT-subgroup 2 showed increases in reactivity to the TMT-paired context, as represented by increased digging behavior during context re-exposure. This is consistent with our hypothesis that higher digging/immobility ratio scores would predict increases in context reactivity. This suggests that defensive digging behavior may be a strong indicator of behavioral reactivity during context re-exposure than immobility. Males, but not females in TMT-subgroup 1 showed avoidance behavior during context re-exposure, also indicative of context reactivity. Interestingly, females in TMT-subgroup 1 showed no behavioral changes during context re-exposure, but were the only group to show increased corticosterone levels during context re-exposure suggesting this subset of female rats exhibited a high stress response to the context. One interpretation of this elevation in corticosterone could be associated with the lack of digging behavior during TMT exposure, such that this group did not engage in proactive coping behaviors (i.e., digging), thus when re-exposed to the TMT-paired context, these rats did not engage in adaptive responses to the stressful context (i.e., digging), resulting in higher HPA-axis activation during context re-exposure. This data pattern suggests a physiological reactivity to the context, which needs to further exploration. Collectively, these data indicate that TMT can produce contextual conditioning, which is consistent with previous work [22], as evidenced by rodents engaging in similar stress-reactive behaviors as during the initial stressor exposure.

Increased startle response during the acoustic startle paradigm is an indicator of hyperarousal behavior, which is a core symptom profile of PTSD [1]. Animal models of traumatic stress, including TMT [47], can trigger increases in hyperarousal responses. We hypothesized that rats with higher digging/immobility ratio scores, would show anxiety-like and hyperarousal behavior post-TMT exposure. Contrary to our hypothesis, only females in TMT subgroup-1 (lower digging/immobility ratio) showed increased startle response (i.e., hyperarousal). For the females, one interpretation of these data is that recruitment of coping strategies such as defensive digging protects against lasting behavioral responsivity in hyperarousal and anxiety-like tests. While a previous study has reported females show hypo-responsiveness of the acoustic startle response following traumatic stress exposure [48] and predator odor stress [25], it is possible that by providing female rats the opportunity to engage in different behavioral coping strategies (i.e., bedding for digging), we were able to separate specific type of responders that represent susceptibility or resiliency to developing lasting behavioral changes in arousal. Conversely, the lack of change in hyperarousal in male rats exposed to TMT remains unclear, but future work could manipulate different startle intensity or examine prepulse inhibition to further dissect this behavioral difference. Additionally, there was no change in approach/avoidance behavior (e.g., anxiety-like behavior) in the light/dark test and zero maze in males or females following TMT exposure. These tests occurred 7-9 days post-TMT exposure, which is consistent with timing from previous studies that show elevated anxiety-like and hyperarousal behavior following predator odor stress (including TMT) that persist over extended periods [22, 24, 49]. However, these previous studies used Sprague-Dawley rats in the elevated plus maze, while the current study assessed anxiety-like behavioral tests using light/dark test and elevate zero maze in Long-Evans male and female rats. Therefore, it is possible the lack of effect in anxiety-like behavior is due to differences in strain and testing paradigms. To address these potential differences, future studies could examine anxiety-like and hyperarousal behavior using different behavioral tests (e.g., social interaction, prepulse inhibition test) and different time points following TMT exposure.

In the current work, we sought to determine whether the stress reactivity during TMT exposure as determined by the TMT subgroups could predict increases in alcohol drinking. Therefore, we hypothesized that differential stress reactivity would predict subsequent increases in alcohol self-administration. First, we observed increases in drinking in female rats in TMT-2 (high digging/low immobility), but not TMT-1 (low digging/high immobility), suggesting stress-reactive behaviors used to classify female rats into TMT-subgroups predicts increases in alcohol self-administration. Second, these increases in alcohol self-administration began to emerge 24 days (12 sessions) after exposure to TMT, suggesting a delay in the escalation of alcohol drinking after stress exposure. This is in line with other studies that have shown increases and long-term (1-3 weeks) persistence in alcohol drinking after exposure to different models of predator odor stress [13, 15, 16], and other work from our lab showing changes in gene expression that emerge 4 weeks following TMT exposure [27]. Third, males in TMT-1 and 2 did not show changes in alcohol self-administration, suggesting that stress reactivity is a more efficient predictor of changes in self-administration in female rats. While there is existing literature on the impact of predator stress and predator odor stress on alcohol drinking in male and female rodents, the results vary between species, strain and stress parameters. For example, a single exposure to dirty rat bedding in mice increased female drinking during access to 2-bottle choice immediately following stressor exposure but suppressed consumption in males [50], while repeated exposure only produced an increase in alcohol intake in a subset of female mice regardless of prior alcohol history [17]. Therefore, this study adds to the limited preclinical data on differences in stress-alcohol interactions by examining individual differences in TMT-induced long-term alcohol consumption in an operant conditioning model in both male and female Long-Evans rats.

Examining how individual differences in response to stress alters alcohol self-administration can lead to a better understanding of behavioral adaptations that persist to modulate long lasting increases in alcohol consumption. Previous studies have examined individual patterns in response to stress and how these differences may potentiate alcohol drinking [13, 15, 16]. However, only a limited number of studies [14] have examined these patterns in both males and females. Here, we show female rats that engaged in more active behavioral responses to TMT compared to passive responses (high digging/low immobility) showed the escalations in alcohol self-administration. Males, regardless of TMT-subgroup engaged in a higher proportion of immobility behavior and did not show any changes in alcohol self-administration. Similarly, female rats in TMT-subgroup 1 that also had high levels of immobility did not show increases in alcohol consumption. These data patterns suggest that immobility behavior during TMT exposure may act as a protective behavioral coping strategy to prevent increases in alcohol drinking after stress exposure.

There are some potential neurobiological mechanisms that could explain why a specific subset of female rats showed increases in alcohol self-administration after stress. First, the mechanisms by which stress impacts alcohol drinking may be different between male and female rats. For example, female, but not male rats show greater ethanol-induced inhibition on action potential firing in basolateral amygdala (BLA) neurons following exposure to the single prolonged stress (SPS) model, suggesting that ethanol plays a larger role in modulating stress-induced excitability in females [51]. In addition, ethanol selectively decreased the amplitude of hyperpolarization-activated current in BLA neurons in SPS-treated males, but not SPS-treated females, further suggesting sex differences in physiological mechanisms that regulate neuronal excitability in response to stress and ethanol [51]. In another study, chronic mild stress increased alcohol consumption and preference in female Wistar rats compared to males, as well as an increase in synaptophysin expression in the frontal cortex, suggestive of elevated presynaptic plasticity in females [52]. In addition, female rats that show an increased in ethanol intake after exposed to dirty rat bedding [17], also show an increase in p450CC (PFC and hippocampus), GABA_A_R_α_2 (PFC-only)CC and synaptophysin (hippocampus-only), all of which play a potential role in addictive behaviors and behavioral responses to stress [53]. While the current study focused on the relationship between individual variability to stress impacts alcohol consumption between males and females, future studies will focus on understanding the neurobiological changes in male and female brains that occur after stressor exposure and how these changes could elucidate potential mechanisms associated with resiliency and susceptible populations to stress.

In conclusion, the current study showed that a rodent model of inescapable, uncontrollable predator odor stress (TMT), produced individual differences in stress-reactive behaviors. Importantly, we show digging behavior during TMT predicts context reactivity (digging) in males and females, while immobility behavior during TMT in females predicts elevated corticosterone levels after context re-exposure and hyperarousal behavior. In addition, specific stress-reactive behaviors including high defensive digging and low immobility behavior used to classify female rats into TMT-subgroups predicted increases in alcohol self-administration, but this was not observed in males. Together, these data suggest that stress-reactive behaviors during predator odor stressor exposure using TMT can elucidate specific behavioral phenotypes in male and female rats offering insight into individual resilience/susceptibility to stress. In addition, these results suggest lasting consequences of TMT on alcohol self-administration in high stress-reactive females, which can help to further understand the impact of stress on escalations in drinking.

## CRediT authorship contribution statement

**Laura C. Ornelas:** Conceptualization, Data curation, Formal analysis, Investigation, Methodology, Validation, Visualization, Roles/Writing – original draft, Writing – review & editing. **Ryan E. Tyler**. Conceptualization, Data curation, Methodology, Writing - review & editing. **Preethi Irukulapati**. Data curation. **Sudheesha Paladugu**. Data curation. **Joyce Besheer**. Conceptualization, Data curation, Formal analysis, Funding acquisition, Investigation, Methodology, Project administration, Supervision, Validation, Visualization, Roles/Writing - original draft, Writing - review & editing.

## Acknowledgement

This work was supported in part by the National Institute of Health AA026537 (JB) and by the Bowles Center for Alcohol Studies. LCO was supported by Diversity Supplement to AA026537. RET was supported by NS007431. The authors thank Jiaqi Liu, Benjamin Weinberg and Abigail Garcia-Baza for their help with behavioral analysis.

## Data Availability Statement

Data available on request from the authors.

## References

1. Association, A.P. Diagnostic and statistical manual of mental disorders 5th ed. 2013, Washington, DC.

2. Kessler, R.C., Berglund, P., Demler, O., Jin, R., Merikangas, K.R., and Walters, E.E. Lifetime prevalence and age-of-onset distributions of DSM-IV disorders in the National Comorbidity Survey Replication. Arch Gen Psychiatry, 2005. 62(6): p. 593–602.

3. Kilpatrick, D.G., Resnick, H.S., Milanak, M.E., Miller, M.W., Keyes, K.M., and Friedman, M.J. National estimates of exposure to traumatic events and PTSD prevalence using DSM-IV and DSM-5 criteria. J Trauma Stress, 2013. 26(5): p. 537–47.

4. Kessler, R.C., Crum, R.M., Warner, L.A., Nelson, C.B., Schulenberg, J., and Anthony, J.C. Lifetime co-occurrence of DSM-III-R alcohol abuse and dependence with other psychiatric disorders in the National Comorbidity Survey. Arch Gen Psychiatry, 1997. 54(4): p. 313–21.

5. Sonne, S.C., Back, S.E., Diaz Zuniga, C., Randall, C.L., and Brady, K.T. Gender differences in individuals with comorbid alcohol dependence and post-traumatic stress disorder. Am J Addict, 2003. 12(5): p. 412–23.

6. Lehavot, K., Stappenbeck, C.A., Luterek, J.A., Kaysen, D., and Simpson, T.L. Gender differences in relationships among PTSD severity, drinking motives, and alcohol use in a comorbid alcohol dependence and PTSD sample. Psychol Addict Behav, 2014. 28(1): p. 42–52.

7. Ralevski, E., Southwick, S., and Petrakis, I. Trauma- and Stress-Induced Craving for Alcohol in Individuals Without PTSD. Alcohol Alcohol, 2020. 55(1): p. 37–43.

8. Connor, K.M. and Davidson, J.R. Development of a new resilience scale: the Connor-Davidson Resilience Scale (CD-RISC). Depress Anxiety, 2003. 18(2): p. 76–82.

9. Russo, S.J., Murrough, J.W., Han, M.H., Charney, D.S., and Nestler, E.J. Neurobiology of resilience. Nat Neurosci, 2012. 15(11): p. 1475–84.

10. Ehlers, A. and Clark, D.M. A cognitive model of posttraumatic stress disorder. Behav Res Ther, 2000. 38(4): p. 319–45.

11. Charney, D.S. Psychobiological mechanisms of resilience and vulnerability: implications for successful adaptation to extreme stress. Am J Psychiatry, 2004. 161(2): p. 195–216.

12. Yehuda, R. and LeDoux, J. Response variation following trauma: a translational neuroscience approach to understanding PTSD. Neuron, 2007. 56(1): p. 19–32.

13. Edwards, S., Baynes, B.B., Carmichael, C.Y., Zamora-Martinez, E.R., Barrus, M., Koob, G.F., and Gilpin, N.W. Traumatic stress reactivity promotes excessive alcohol drinking and alters the balance of prefrontal cortex-amygdala activity. Transl Psychiatry, 2013. 3: p. e296.

14. Albrechet-Souza, L., Schratz, C.L., and Gilpin, N.W. Sex differences in traumatic stress reactivity in rats with and without a history of alcohol drinking. Biol Sex Differ, 2020. 11(1): p. 27.

15. Manjoch, H., Vainer, E., Matar, M., Ifergane, G., Zohar, J., Kaplan, Z., and Cohen, H. Predator-scent stress, ethanol consumption and the opioid system in an animal model of PTSD. Behav Brain Res, 2016. 306: p. 91–105.

16. Weera, M.M., Schreiber, A.L., Avegno, E.M., and Gilpin, N.W. The role of central amygdala corticotropin-releasing factor in predator odor stress-induced avoidance behavior and escalated alcohol drinking in rats. Neuropharmacology, 2020. 166: p. 107979.

17. Finn, D.A., Helms, M.L., Nipper, M.A., Cohen, A., Jensen, J.P., and Devaud, L.L. Sex differences in the synergistic effect of prior binge drinking and traumatic stress on subsequent ethanol intake and neurochemical responses in adult C57BL/6J mice. Alcohol, 2018. 71: p. 33–45.

18. King, C.E. and Becker, H.C. Oxytocin attenuates stress-induced reinstatement of alcohol seeking behavior in male and female mice. Psychopharmacology (Berl), 2019. 236(9): p. 2613–2622.

19. Dopfel, D., Perez, P.D., Verbitsky, A., Bravo-Rivera, H., Ma, Y., Quirk, G.J., and Zhang, N. Individual variability in behavior and functional networks predicts vulnerability using an animal model of PTSD. Nat Commun, 2019. 10(1): p. 2372.

20. Cohen, H. and Zohar, J. An animal model of posttraumatic stress disorder: the use of cut-off behavioral criteria. Ann N Y Acad Sci, 2004. 1032: p. 167–78.

21. Shallcross, J., Hamor, P., Bechard, A.R., Romano, M., Knackstedt, L., and Schwendt, M. The Divergent Effects of CDPPB and Cannabidiol on Fear Extinction and Anxiety in a Predator Scent Stress Model of PTSD in Rats. Front Behav Neurosci, 2019. 13: p. 91.

22. Schwendt, M., Shallcross, J., Hadad, N.A., Namba, M.D., Hiller, H., Wu, L., Krause, E.G., and Knackstedt, L.A. A novel rat model of comorbid PTSD and addiction reveals intersections between stress susceptibility and enhanced cocaine seeking with a role for mGlu5 receptors. Transl Psychiatry, 2018. 8(1): p. 209.

23. Brodnik, Z.D., Black, E.M., and Espana, R.A. Accelerated development of cocaine-associated dopamine transients and cocaine use vulnerability following traumatic stress. Neuropsychopharmacology, 2020. 45(3): p. 472–481.

24. Brodnik, Z.D., Black, E.M., Clark, M.J., Kornsey, K.N., Snyder, N.W., and Espana, R.A. Susceptibility to traumatic stress sensitizes the dopaminergic response to cocaine and increases motivation for cocaine. Neuropharmacology, 2017. 125: p. 295–307.

25. Albrechet-Souza, L. and Gilpin, N.W. The predator odor avoidance model of post-traumatic stress disorder in rats. Behav Pharmacol, 2019. 30(2 and 3-Spec Issue): p. 105–114.

26. Rosen, J.B., Asok, A., and Chakraborty, T. The smell of fear: innate threat of 2,5-dihydro-2,4,5-trimethylthiazoline, a single molecule component of a predator odor. Front Neurosci, 2015. 9: p. 292.

27. Tyler, R.E., Weinberg, B., Lovelock, D., Ornelas, L.C., and Besheer, J. Exposure to the predator odor TMT induces early and late differential gene expression related to stress and excitatory synaptic function throughout the brain in male rats. Genes, Brain and Behavior, 2020.

28. Verbitsky, A., Dopfel, D., and Zhang, N. Rodent models of post-traumatic stress disorder: behavioral assessment. Transl Psychiatry, 2020. 10(1): p. 132.

29. De Boer, S.F. and Koolhaas, J.M. Defensive burying in rodents: ethology, neurobiology and psychopharmacology. Eur J Pharmacol, 2003. 463(1-3): p. 145–61.

30. Arakawa, H. Ontogeny of sex differences in defensive burying behavior in rats: effect of social isolation. Aggress Behav, 2007. 33(1): p. 38–47.

31. Fucich, E.A. and Morilak, D.A. Shock-probe Defensive Burying Test to Measure Active versus Passive Coping Style in Response to an Aversive Stimulus in Rats. Bio Protoc, 2018. 8(17).

32. Riittinen, M.L., Lindroos, F., Kimanen, A., Pieninkeroinen, E., Pieninkeroinen, I., Sippola, J., Veilahti, J., Bergstrom, M., and Johansson, G. Impoverished rearing conditions increase stress-induced irritability in mice. Dev Psychobiol, 1986. 19(2): p. 105–11.

33. Neal, S., Kent, M., Bardi, M., and Lambert, K.G. Enriched Environment Exposure Enhances Social Interactions and Oxytocin Responsiveness in Male Long-Evans Rats. Front Behav Neurosci, 2018. 12: p. 198.

34. Wallace, K.J. and Rosen, J.B. Predator odor as an unconditioned fear stimulus in rats: elicitation of freezing by trimethylthiazoline, a component of fox feces. Behav Neurosci, 2000. 114(5): p. 912–22.

35. Homiack, D., O’Cinneide, E., Hajmurad, S., Dohanich, G.P., and Schrader, L.A. Effect of acute alarm odor exposure and biological sex on generalized avoidance and glutamatergic signaling in the hippocampus of Wistar rats. Stress, 2018. 21(4): p. 292–303.

36. Homiack, D., O’Cinneide, E., Hajmurad, S., Barrileaux, B., Stanley, M., Kreutz, M.R., and Schrader, L.A. Predator odor evokes sex-independent stress responses in male and female Wistar rats and reduces phosphorylation of cyclic-adenosine monophosphate response element binding protein in the male, but not the female hippocampus. Hippocampus, 2017. 27(9): p. 1016–1029.

37. Metna-Laurent, M., Soria-Gomez, E., Verrier, D., Conforzi, M., Jego, P., Lafenetre, P., and Marsicano, G. Bimodal control of fear-coping strategies by CB(1) cannabinoid receptors. J Neurosci, 2012. 32(21): p. 7109–18.

38. Koolhaas, J.M., Korte, S.M., De Boer, S.F., Van Der Vegt, B.J., Van Reenen, C.G., Hopster, H., De Jong, I.C., Ruis, M.A., and Blokhuis, H.J. Coping styles in animals: current status in behavior and stress-physiology. Neurosci Biobehav Rev, 1999. 23(7): p. 925–35.

39. Makhijani, V.H., Van Voorhies, K., and Besheer, J. The mineralocorticoid receptor antagonist spironolactone reduces alcohol self-administration in female and male rats. Pharmacol Biochem Behav, 2018. 175: p. 10–18.

40. Asok, A., Ayers, L.W., Awoyemi, B., Schulkin, J., and Rosen, J.B. Immediate early gene and neuropeptide expression following exposure to the predator odor 2,5-dihydro-2,4,5-trimethylthiazoline (TMT). Behav Brain Res, 2013. 248: p. 85–93.

41. Hwa, L.S., Neira, S., Pina, M.M., Pati, D., Calloway, R., and Kash, T.L. Predator odor increases avoidance and glutamatergic synaptic transmission in the prelimbic cortex via corticotropin-releasing factor receptor 1 signaling. Neuropsychopharmacology, 2019. 44(4): p. 766–775.

42. Falconer, E.M. and Galea, L.A. Sex differences in cell proliferation, cell death and defensive behavior following acute predator odor stress in adult rats. Brain Res, 2003. 975(1-2): p. 22–36.

43. Fucich, E.A., Paredes, D., and Morilak, D.A. Therapeutic Effects of Extinction Learning as a Model of Exposure Therapy in Rats. Neuropsychopharmacology, 2016. 41(13): p. 3092–3102.

44. Gruene, T.M., Flick, K., Stefano, A., Shea, S.D., and Shansky, R.M. Sexually divergent expression of active and passive conditioned fear responses in rats. Elife, 2015. 4.

45. Shansky, R.M. Sex differences in PTSD resilience and susceptibility: Challenges for animal models of fear learning. Neurobiol Stress, 2015. 1: p. 60–65.

46. Colom-Lapetina, J., Li, A.J., Pelegrina-Perez, T.C., and Shansky, R.M. Behavioral Diversity Across Classic Rodent Models Is Sex-Dependent. Front Behav Neurosci, 2019. 13: p. 45.

47. Hebb, A.L., Zacharko, R.M., Gauthier, M., and Drolet, G. Exposure of mice to a predator odor increases acoustic startle but does not disrupt the rewarding properties of VTA intracranial self-stimulation. Brain Res, 2003. 982(2): p. 195–210.

48. Pooley, A.E., Benjamin, R.C., Sreedhar, S., Eagle, A.L., Robison, A.J., Mazei-Robison, M.S., Breedlove, S.M., and Jordan, C.L. Sex differences in the traumatic stress response: PTSD symptoms in women recapitulated in female rats. Biol Sex Differ, 2018. 9(1): p. 31.

49. Lim, J., Igarashi, M., Jung, K.M., Butini, S., Campiani, G., and Piomelli, D. Endocannabinoid Modulation of Predator Stress-Induced Long-Term Anxiety in Rats. Neuropsychopharmacology, 2016. 41(5): p. 1329–39.

50. Cozzoli, D.K., Tanchuck-Nipper, M.A., Kaufman, M.N., Horowitz, C.B., and Finn, D.A. Environmental stressors influence limited-access ethanol consumption by C57BL/6J mice in a sex-dependent manner. Alcohol, 2014. 48(8): p. 741–54.

51. Ornelas, L.C. and Keele, N.B. Sex Differences in the Physiological Response to Ethanol of Rat Basolateral Amygdala Neurons Following Single-Prolonged Stress. Front Cell Neurosci, 2018. 12: p. 219.

52. Marco, E.M., Ballesta, J.A., Irala, C., Hernandez, M.D., Serrano, M.E., Mela, V., Lopez-Gallardo, M., and Viveros, M.P. Sex-dependent influence of chronic mild stress (CMS) on voluntary alcohol consumption; study of neurobiological consequences. Pharmacol Biochem Behav, 2017. 152: p. 68–80.

53. Devaud, L.L., Alavi, M., Jensen, J.P., Helms, M.L., Nipper, M.A., and Finn, D.A. Sexually divergent changes in select brain proteins and neurosteroid levels after a history of ethanol drinking and intermittent PTSD-like stress exposure in adult C57BL/6J mice. Alcohol, 2020. 83: p. 115–125.

